# A data-driven, meaningful, easy to interpret, population-independent accelerometer outcome variable for global surveillance

**DOI:** 10.1101/604694

**Authors:** Alex V. Rowlands, Lauren B. Sherar, Stuart J. Fairclough, Tom Yates, Charlotte L. Edwardson, Deirdre M. Harrington, Melanie J. Davies, Fehmidah Munir, Kamlesh Khunti, Victoria H. Stiles

**Affiliations:** Diabetes Research Centre, University of Leicester, Leicester General Hospital, Leicester, UK; NIHR Leicester Biomedical Research Centre, UK; Alliance for Research in Exercise, Nutrition and Activity (ARENA), Sansom Institute for Health Research, Division of Health Sciences, University of South Australia, Adelaide, Australia; School of Sport, Exercise and Health Sciences, Loughborough University, Loughborough, UK; Movement Behaviours, Health, and Wellbeing Research Group, Department of Sport and Physical Activity, Edge Hill University, Ormskirk, UK; NIHR Collaboration for Leadership in Applied Health Research and Care East Midlands, Leicester General Hospital, UK; Sport and Health Sciences, College of Life and Environmental Sciences, University of Exeter, Exeter, UK

**Keywords:** MVPA, public health guidelines, population-independent, accelerometer metrics, GENEActiv, Axivity, ActiGraph

## Abstract

**Background:** Accelerometer-driven physical activity guidelines are not available, likely due to the lack of consensus on meaningful and interpretable accelerometer outcomes. The aim of this paper is to demonstrate how a data-driven accelerometer metric, the acceleration above which a person’s most active minutes are accumulated, can a) quantify the prevalence of meeting current physical activity guidelines for global surveillance and b) moving forward, could inform accelerometer-driven physical activity guidelines. Unlike cut-point methods, the metric is population-independent (e.g. age) and comparable across datasets.

**Methods:** Secondary data analyses were carried out on five datasets using wrist-worn accelerometers: children (N=145), adolescent girls (N=1669), office workers (N=114), pre- (N=1218) and post- (N=1316) menopausal women, and adults with type 2 diabetes (N=475). Open-source software (GGIR) was used to generate the magnitude of acceleration above which a person’s most active 60, 30 and 2 minutes are accumulated: M60_ACC_; M30_ACC_ and M2_ACC_, respectively.

**Results:** The proportion of participants with M60_ACC_ (children) and M30_ACC_ (adults) values higher than accelerations indicative of brisk walking (i.e., moderate-to-vigorous physical activity) ranged from 17-68% in children and 15%-81% in adults, tending to decline with age. The proportion of pre-and post-menopausal women with M2_ACC_ values indicative of running and thus meeting recently presented thresholds for bone health ranged from 6-13%.

**Conclusion:** These metrics can be used for global surveillance of physical activity, including assessing prevalence of meeting the current physical activity guidelines, across the lifespan. Translation of acceleration magnitudes into indicative activities provides a public health friendly interpretation of results. As accelerometer and corresponding health data accumulate it will be possible to interpret the metrics relative to age- and sex-specific norms and derive evidence-based physical activity guidelines directly from accelerometer data for use in future global surveillance. This is where the key advantages of these metrics lie.

## Introduction

National and/or large-scale surveys of physical activity through accelerometers are now commonplace in many countries worldwide^1-5^. The World Health Organisation’s recent Global Activity Action Plan on Physical Activity 2018-2030^6^ highlights monitoring and surveillance, using robust and reliable data, as the cornerstone to the implementation and evaluation of national strategies. Accelerometers provide a valid measure of physical activity^7^; however, a lack of consensus on robust and consistent methods to reduce and analyse data to create meaningful and easy to interpret outcome variables, is hampering monitoring and evaluation activities.

For example, epidemiological studies and surveillance studies frequently create variables from accelerometer-assessed moderate-to-vigorous physical activity (MVPA) using intensity cut-points. The problems with using cut-points to quantify activity are well documented but, briefly, include: (1) cut-points are protocol-, and population- (e.g. age-group) specific, leading to results that are not comparable across studies^8-10^; (2) two participants with similar levels of activity score very differently if one has activity falling just above the cut-point and one has activity falling just below the cut-point; (3) many participants fail to obtain any activity above cut-points (particularly in the vigorous range), consequently a large number of people simply score zero minutes. Recently, in an examination of how cut-points influence estimates of physical activity, Migueles et al.^11,p1^ stated that it was ‘not possible (and probably will never be) to know the prevalence of meeting the physical activity guidelines based on accelerometer data’. Clearly a new approach to analysing and interpreting accelerometer data is needed.

An alternative approach is to identify the minimum acceleration value above which a person’s most active minutes, for example 30 mins (M30_ACC_), is accumulated. The active minutes can be accumulated in any way across the day, with no need for the activity to be in bouts, in line with recent physical activity recommendations^12^. With this approach the metric is population-independent and derived from directly measured acceleration, thus not relying on assumptions as cut-points do^9^, and the intensity is captured regardless of level of activity with no person scoring zero. This bears similarities to the peak 30 min walking cadence (steps/min) proposed by Tudor-Locke and colleagues^13^ as a practical estimate of activity intensity.

Moving forward, as accelerometer and corresponding health data accumulate, these data-driven population-independent metrics could be used to inform accelerometer-driven physical activity guidelines as recommended by Troiano et al.^10^, rather than inappropriately evaluating physical activity assessed by accelerometer cut-points to guidelines developed from self-report data, which are conceptually different^10^. For example, the M30_ACC_ and/or M60_ACC_ that is positively associated with a given health marker, e.g. adiposity, could be determined. This M30_ACC_ and/or M60_ACC_ value could then be used for surveillance which, importantly, would facilitate surveillance using the same physical activity metric as used to garner the evidence. As data accumulate, it would be possible to interpret the M30_ACC_ and M60_ACC_ relative to age- and sex-specific norms and/or relative to values associated with health markers.

To facilitate public-health recommendations, translation of the metrics to public-health friendly indicative activity types is desirable, e.g. brisk walking, and/or MVPA. This translation is necessarily population-specific and thus bears similarities to cut-point analyses. However, crucially this is <u>only</u> in the translation of the data for activity recommendations because all analyses are carried out on the population-independent metrics^9^. In contrast, when using cut-points, thresholds are imposed on the data from the outset to collapse data into categories for analysis, rendering it impossible to subsequently compare any datasets deploying different cut-points.

For example, assume that a child has an M60_ACC_ of 225 m*g*. Until we have the data to compare this to accelerometer-driven physical activity guidelines, we can assess whether the child is meeting the current 60 min daily MVPA guideline^12^ by comparing the M60_ACC_ value to indicative activities, or the various cut-points available. According to the 200 m*g* MVPA cut-point (indicative of brisk walking) published by Hildebrand et al.^14^, the child exceeds the 60 minutes of MVPA per day recommendations^12^, while according to a more stringent 250 m*g* MVPA cut-point published by Phillips et al.^15^, the child does not quite reach the recommendations. If a cut-point approach had been used to analyse the data, the child’s score could not be compared to any alternative cut-point or threshold.

For the purposes of a simple demonstration of how these metrics could be used for surveillance of adherence to current physical activity guidelines^12^, we looked at the daily average acceleration above which the most active 30 mins (M30_ACC_, adults) or 60 mins (M60_ACC_, children) was obtained. It would be possible to alter the number of minutes over which the minimum acceleration is considered, depending on the health outcome of interest or the guideline being assessed. For example, in a large cross-sectional observational study, Stiles et al.^16^ demonstrated that accumulating 1-2 minutes of accelerometer-assessed high intensity activity, equivalent to running, was associated with bone health in pre- and post-menopausal women.

The primary aim of this paper is to demonstrate how the acceleration above which a person’s most active minutes are accumulated, can be used to quantify prevalence of meeting existing physical activity guidelines. A secondary aim is to illustrate how in the future, as accelerometer and corresponding health data accumulate, these population-independent metrics could be used to inform accelerometer-driven physical activity guidelines, which is where the key advantages of these metrics lie.

## Methods

Secondary data analyses were carried out on five diverse datasets: 10 y old children^17^, adolescent girls^18,19^, adult office workers^20^, pre- and post-menopausal women^16^, and adults with type 2 diabetes. All participants gave assent (children and adolescent girls) or informed consent (adults). Parents/guardians of the children gave written informed consent and parents/guardians of the adolescent girls returned an opt-out consent form if they did not want their child to participate. All studies received the appropriate institutional ethics approval.

In all samples, wrist worn accelerometers were worn 24 h a day for up to 7-days. The children and adult office workers wore the ActiGraph GT9X (ActiGraph, Pensacola, FL, USA), the adolescent girls and the adults with type 2 diabetes wore the GENEActiv (ActivInsights Ltd, Cambridgeshire, UK) and the pre- and post-menopausal women wore the Axivity AX3 (Axivity, Newcastle, UK). The pre- and post-menopausal women wore the monitor on their dominant wrist, all other samples wore monitors on the non-dominant wrist. All monitors were initialised to record accelerations at 100 Hz, except the adult office workers whose monitors were initialised at 30 Hz.

ActiGraphs were initialised and downloaded using ActiLife version 6.11.9 (ActiGraph, Pensacola, FL, USA). Data were saved in raw format as GT3X files, before being converted to raw csv file format for signal processing. GENEActivs were initialised and data downloaded in binary format using GENEActiv PC (version 3.1). Axivity data were downloaded from UK Biobank in .cwa format, auto-calibrated, resampled (100 Hz) and converted to .wav format using open-source software (Omgui Version 1.0.0.28; Open Movement, Newcastle, UK).

All accelerometer files were processed and analysed with R-package GGIR version 1.6-7 (http://cran.r-project.org)^21,22^. Signal processing in GGIR included auto-calibration using local gravity as a reference^21^ (apart from the Axivity files which were auto-calibrated when converted to .wav files); detection of sustained abnormally high values; detection of non-wear; and calculation of the average magnitude of dynamic acceleration corrected for gravity (Euclidean Norm minus 1 g, ENMO). These were averaged over 1 or 5 s epochs (1s: pre- and post-menopausal women (UK Biobank); 5 s: children, adolescent girls, adult office workers and adults with type 2 diabetes) and expressed in milli-gravitational units (m*g*).

Participants were excluded if their accelerometer files showed: post-calibration error greater than 0.01 *g* (10 m*g*), fewer than three days of valid wear (defined as >16 h per day), or wear data wasn’t present for each 15 min period of the 24 h cycle. The following metrics were generated and averaged across all valid days: average acceleration; intensity gradient (intensity distribution^23^); acceleration above which a person’s most active X minutes (MX_ACC_) are accumulated: M60_ACC_ (m*g*); M30_ACC_ (m*g*), M2_ACC_ (m*g*), (GGIR qlevels (0,24 hours): 1380/1440, 1410/1440 and 1438/1440). As acceleration measured at the dominant wrist is approximately 10% higher than the non-dominant^24^, magnitudes of M60_ACC_, M30_ACC_ and M2_ACC_ were reduced by 10% for dominant wrist placement (pre- and post-menopausal women).

### Analyses

Descriptive statistics were calculated using mean (standard deviation (SD)) for continuous variables and percentage for categorical variables.

Percentiles (5^th^ - 95^th^ percentile) were graphed for females (all samples) and males (where available) for the M60_ACC_, M30_ACC_ and M2_ACC_. Presenting percentiles for each metric illustrates the magnitude of the most active X minutes, from the least to the most active participants, within each sample. To address our primary aim, the proportion of each sample meeting the MVPA physical activity guidelines, operationalised for the purposes of this demonstration as a daily average of 30 min for adults and 60 mins for children and adolescents, was calculated. For MVPA, we used acceleration values indicative of a brisk walk (5 km/h, ≅ 3.6 METs: 170 m*g* adults; 200 m*g* children^14^) and of a fast walk (5.6 km/h, ≅ 4.5 METs: 250 m*g* adults; 300 m*g* children^14,25,26^). In addition, the proportion of pre- and post-menopausal women meeting the recently proposed accelerometer-driven guide of 2 min high-intensity activity associated with bone health^16^ was calculated. The thresholds (>1000 m*g* (medium run) pre-menopausal, > 750 m*g*, post-menopausal (slow run)) were generated using dominant wrist data^16^, so are adjusted by −10%^24^.

## Results

Descriptive characteristics are presented in Table 1. Valid accelerometer data files were available for 64% of 10 y old children, 96% of adolescent girls, 78% of adult office workers and 99% of adults with type 2 diabetes. All accelerometer files for the pre- and post-menopausal women from UK Biobank meeting the criteria of Stiles et al.^16^ were available and included (see Stiles et al.^18^ for details).

**Table 1.** Descriptive characteristics of the five datasets.

|  |  | 9-10 y old children | Adolescent girls |  | Adult office workers | Women: UK Biobank |  | Adults with type 2 diabetes |
| --- | --- | --- | --- | --- | --- | --- | --- | --- |
|  |  | (N=145) | 11-12 y (N=974) | 13-14 y (N=695) | (N=114) | Pre-menopausal (N=1218) | Post-menopausal (N=1316) | (N = 475) |
| Sex (%) | Males | 42.8 | 0 | 0 | 20.4 | 0 | 0 | 64 |
|  | Females | 57.2 | 100 | 100 | 79.6 | 100 | 100 | 36 |
| Age (y) |  | 9.6 (0.3) | 12.3 (0.4) | 13.6 (0.4) | 41.2 (10.9) | 46.2 (3.9) | 59.0 (5.1) | 64.2 (8.7) |
| Body size | Height (cm) | 137.5 (5.9) | 153.5 (7.7) | 159.5 (6.8) | 165.9 (7.5) | 164.9 (6.0) | 163.2 (6.1) | 168.6 (11.4) |
|  | Mass (kg) | 35.2 (8.2) | 45.5 (10.8) | 53.6 (12.8) | 73.1 (17.3) | 65.4 (12.0) | 68.1 (11.8) | 107.5 (14.5) |
|  | Body mass index (BMI) (kg.m<sup>-2</sup>) | 18.5 (3.3) | 19.2(3.6) | 20.9 (4.3) | 26.5 (5.9) | 24.9 (4.2) | 25.6 (4.4) | 31.4 (5.4) |
|  | **zBMI | 0.63 (1.19) | 0.08 (1.30) | 0.34 (1.33) |  |  |  |  |
| Accelerometer | Brand | ActiGraph GT9X | GENEActiv | GENEActiv | ActiGraph GT9X | Axivity | Axivity | GENEActiv |
|  | Wrist | Non-dominant | Non-dominant | Non-dominant | Non-dominant | Dominant | Dominant | Non-dominant |
| *Physical activity | Average acceleration (mg) | 45.8 (13.1) | 37.8 (9.0) | 34.3 (7.9) | 26.9 (7.7) | *30.6 (8.5) | *27.1 (7.0) | 22.0 (7.3) |
|  | †Intensity gradient | -1.96 (0.14) | -2.19 (0.15) | -2.28 (0.17) | -2.55 (0.22) | -2.66 (0.16) | -2.74 (0.16) | -2.74 (0.20) |
|  | ‡M60<sub>ACC</sub> (mg) | 216.9 (71.5) | 180.4 (42.9) | 166.7 (37.7) | 129.1 (37.9) | *158.3 (47.7) | *139.1 (34.1) | 103.9 (36.3) |
|  | ‡M30<sub>ACC</sub> (mg) | 363.9 (135.5) | 260.6 (75.8) | 233.0 (63.80) | 188.1 (95.6) | *226.4 (85.9) | *191.7(56.2) | 136.9 (50.5) |
|  | ‡M2<sub>ACC</sub> (mg) | 1545.1 (518.8) | 954.2 (323.7) | 771.9 (301.5) | 426.5 (216.2) | *522.2 (228.9) | *503.8 (160.3) | 305.0 (115.0) |

Values are mean (standard deviation) for continuous variables and % for categorical variables.
\* Reduced by 10% as acceleration measured at the dominant wrist is approximately 10% higher than measured at the non-dominant<sup>24</sup>.
\*\* zBMI: BMI expressed in z-scores for sex and age according to reference curves for the UK<sup>27</sup>.
†Measure of the intensity distribution of the 24 h activity profile, see Rowlands et al.<sup>23</sup>. A more negative gradient reflects a steeper drop with little time accumulated at mid-range and higher intensities, while a less negative gradient reflects a shallower drop with more time spread across the intensity range.
‡M60<sub>ACC</sub>, M30<sub>ACC</sub>, M2<sub>ACC</sub>: acceleration above which a person's most active minutes (X min, MX<sub>ACC</sub>) are accumulated.

Figures 1 and 2 show percentile plots for M60_ACC_, M30_ACC_ and M2_ACC_ for females and males, respectively, in order of increasing sample mean age. Accelerations associated with a brisk walk (5 km/h), fast walk (5.6 km/h) and run (≥8 km/h) are marked on the y-axes to illustrate how the data could be translated in a public-health friendly way^14^. The expected age-related decline in intensity of physical activity was relatively greater the fewer minutes considered (i.e. M2_ACC_ relative to M30_ACC_, and M30_ACC_ relative to M60_ACC_), but also for higher percentiles (i.e. higher intensity) within a given outcome (Figures 1b-c, 2b-c). Sex differences were most evident in 10 y old children, with the intensity of boys’ activity greater than that of girls’ (Figures 1a-c compared to 2a-c).

**Figure 1:**
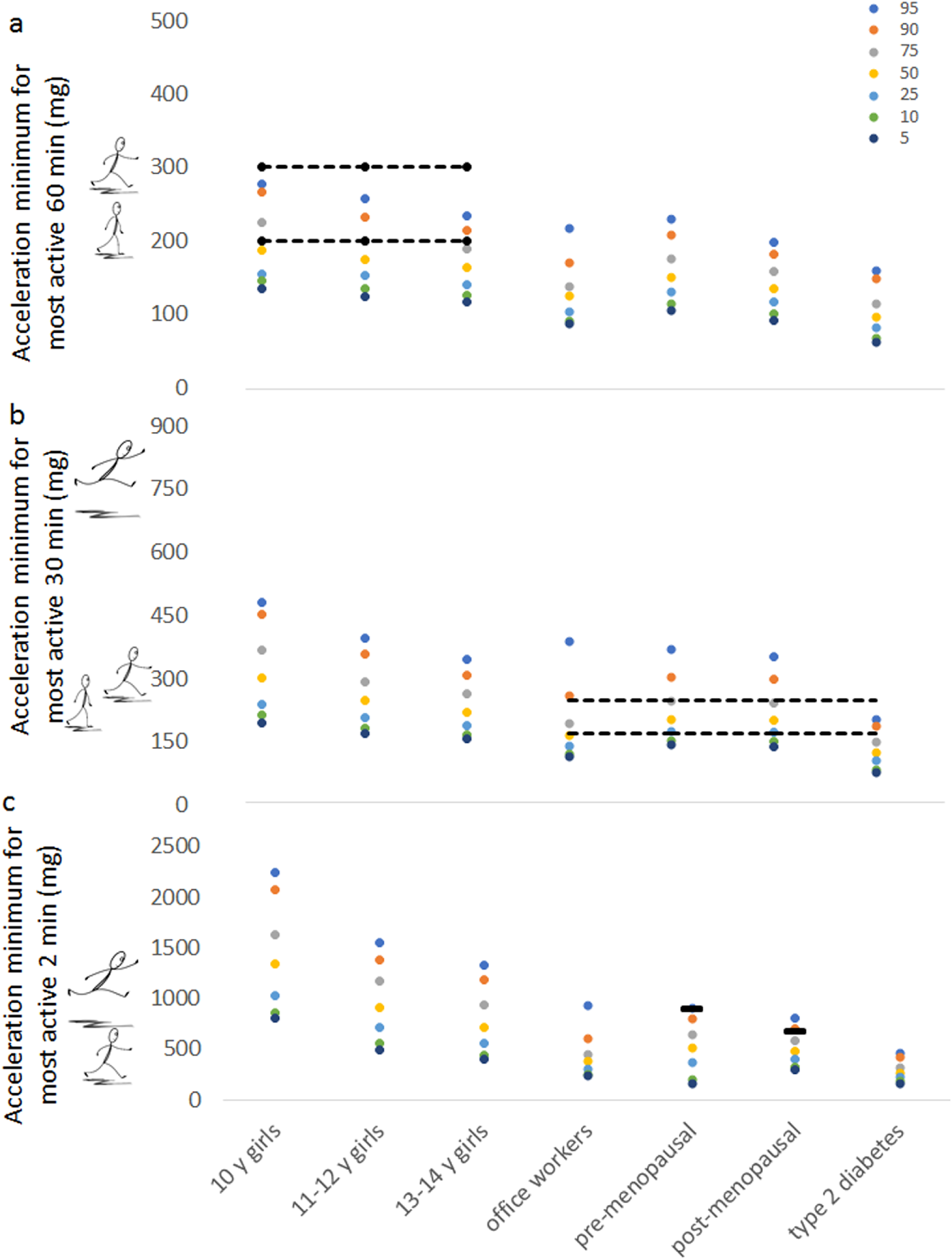
Percentiles for the magnitude of acceleration above which the females’ most active (a) 60, (b) 30 and (c) 2 minutes are accumulated: M60_ACC_; M30_ACC_ and M2_ACC_ (m*g*). Black dashes /dashed lines represent: (a) M60_ACC_ and (b) M30_ACC_ at the intensity of a brisk walk (lower dashed line) or fast walk (upper dashed line); (c) M2_ACC_ at the bone health threshold: medium running for pre-menopausal women and slow running for post-menopausal women^14^

**Figure 2:**
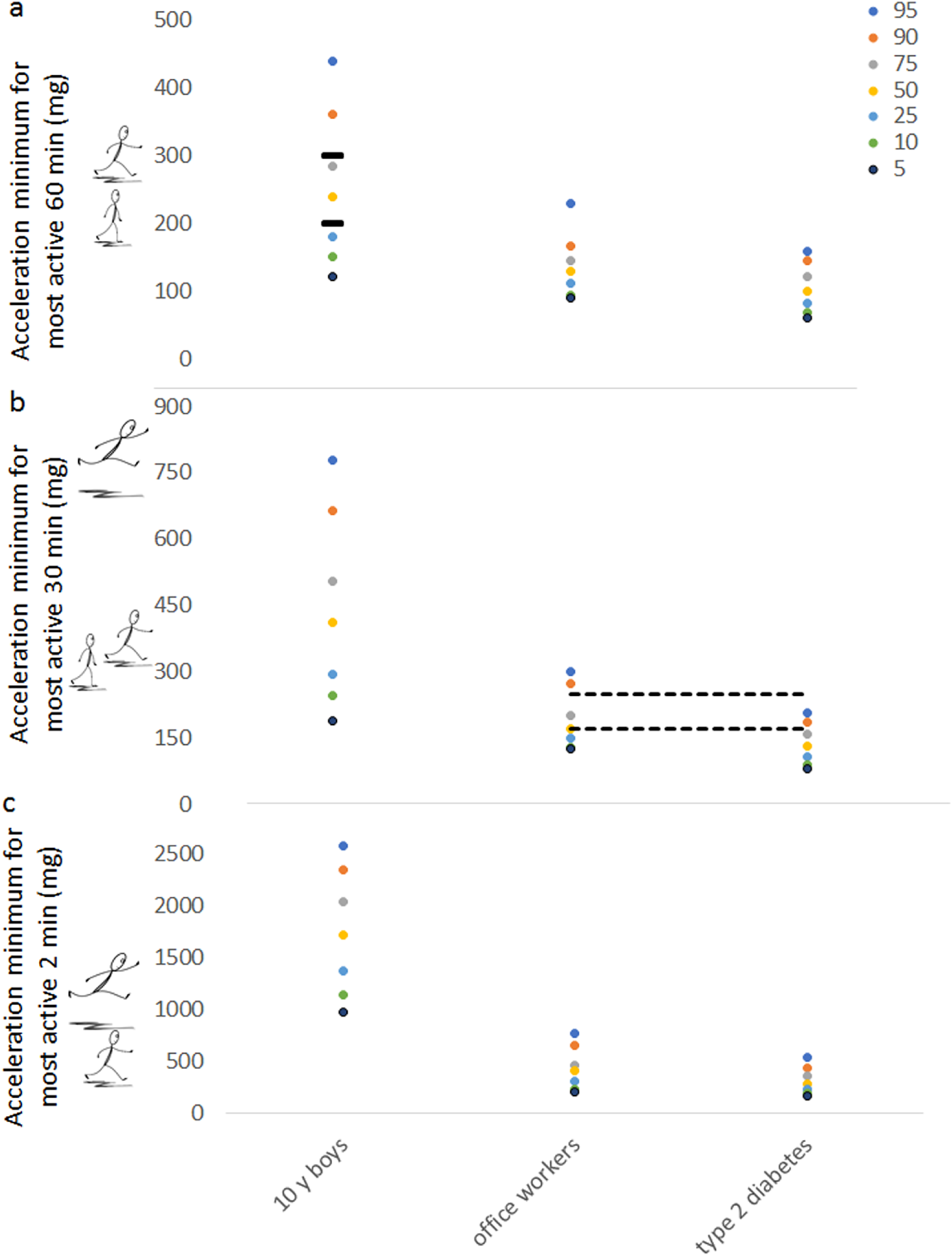
Percentiles for the magnitude of acceleration above which the males’ most active (a) 60, (b) 30 and (c) 2 minutes are accumulated: M60_ACC_; M30_ACC_ and M2_ACC_ (m*g*). Black dashes /dashed lines represent: (a) M60_ACC_ and (b) M30_ACC_ at the intensity of a brisk walk (lower dashes / dashed line) or fast walk (upper dashes / dashed line)

Table 2 shows the proportion of each sample meeting MVPA guidelines operationalised as 60 min per day (children) or 30 min per day (adults) of brisk walking or fast walking. The MX_ACC_ above which the most active time is accumulated is shown for those meeting and not meeting the guidelines. The proportions of pre- and post-menopausal women meeting the recent accelerometer-derived guide proposed for bone health (2 minutes >1000 m*g* (medium run) pre-menopausal, >750 m*g* (slow run) post-menopausal (18)) were 6% and 13%, respectively (Figure 1c).

**Table 2:** Proportion of each sample meeting MVPA guidelines operationalised as 60 min per day (children) or 30 min per day (adults) of brisk walking or fast walking

|  |  | Brisk walk |  | Fast walk |  |
| --- | --- | --- | --- | --- | --- |
|  |  | Female | Male | Female | Male |
| Children | 60 min | >200 mg | >200 mg | >300 mg | >300 mg |
| 10 y olds | % meeting guideline | 39% | 68% | 2% | 21% |
|  | NO: M60<sub>ACC</sub> | 164.9 (21.3) | 157.8 (27) | 192.0 (42.2) | 212.1 (53.1) |
|  | YES: M60<sub>ACC</sub> | 242.3 (28.4) | 289.1 (74.4) | 302.2 (1.64) | 377.2 (70.2) |
| 11-12 y olds | % meeting guideline | 26% |  | 2% |  |
|  | NO: M60<sub>ACC</sub> | 161.4 (23.3) |  | 177.6 (36.8) |  |
|  | YES: M60<sub>ACC</sub> | 235.6 (39.0) |  | 355.2 (43.30) |  |
| 13-14 y olds | % meeting guideline | 17% |  | 1% |  |
|  | NO: M60<sub>ACC</sub> | 154.6 (25.1) |  | 164.4 (25.6) |  |
|  | YES: M60<sub>ACC</sub> | 228.1 (30.1) |  | 320.1 (10.5) |  |
| Adults | 30 min | >170 mg | >170 mg | >250 mg | >250 mg |
| Office workers | % Meeting guideline | 45% | 55% | 11% | 14% |
|  | NO: M30<sub>ACC</sub> | 141.7 (18.2) | 147.7 (16.0) | 161.4 (30.9) | 167.5 (27.4) |
|  | YES: M30<sub>ACC</sub> | 246.8 (133.6) | 212.8 (46.0) | 427.9 (188.5) | 282.8 (19.5) |
| Pre-menopausal women | % meeting guideline | 81% |  | 25% |  |
|  | NO: M30<sub>ACC</sub> | 150.8 (15.1) |  | 191.2 (31.3) |  |
|  | YES: M30<sub>ACC</sub> | 244.5 (86.0) |  | 334.5 (107.6) |  |
| Post-menopausal women | % meeting guideline | 63% |  | 11% |  |
|  | NO: M30<sub>ACC</sub> | 145.6 (17.6) |  | 177.3 (33.5) |  |
|  | YES: M30<sub>ACC</sub> | 219.0 (53.4) |  | 303.4 (69.9) |  |
| Adults with type 2 diabetes | % meeting guideline | 15% | 16% | 2% | 3% |
|  | NO: M30<sub>ACC</sub> | 118.7 (23.9) | 123.9 (26.5) | 129.2 (35.1) | 132.8 (33.9) |
|  | YES: M30<sub>ACC</sub> | 203.8 (28.4) | 221.2 (89.1) | 268.8 (15.4) | 354.5 (0) |

## Discussion

Given the rising use of accelerometers, including their use in large-scale surveys^e.g.1-5^, it is important to have simple to derive and easy to interpret accelerometer variables that can be used to compare physical activity across datasets/populations/countries. This would facilitate global surveillance and the development of evidence-based physical activity guidelines directly from accelerometer data. As data accumulate, physical activity of groups and individuals can be interpreted relative to age- and sex-specific norms and/or relative to values associated with health markers. While the values themselves are not immediately intuitive, this is also true of many metrics that are commonly used by researchers, clinicians and the public^28^. For example, risk thresholds for health markers such as body mass index, blood pressure, and cholesterol are routinely used and widely understood. As outlined by Welk et al.^28^, a range of instruments are used to obtain measures of blood pressure, but the use of a standardised metric makes it possible for researchers, clinicians and patients to discuss a common number. This would also be possible with widespread use of standardised population-independent accelerometer measures of physical activity.

In this paper, we demonstrate how presenting percentiles for population-independent metrics such as the M60_ACC_ and M30_ACC_ can be used now to estimate adherence to current MVPA guidelines. The numerous problems associated with applying cut-points to accelerometer data^8-10^ are avoided as the data and results presented are data-driven. Comparison to any indicative activity, cut-point or, more importantly, any future health-related accelerometer threshold is possible and can be carried out post-hoc with no access to the original data needed. Further, demographic-specific translations can be carried out post-hoc to facilitate public-health friendly recommendations using accelerations representative of typical activities. Crucially, population-specific translation is only for interpretation and has no bearing on analyses or results presented. This means the metrics and results retain their population-independence^9^.

Further, comparison or translation is not tied to an exact acceleration value for an indicative activity or cut-point. For example, if a child accumulates 60 min of activity in the acceleration range of 185 – 199 m*g*, their M60_ACC_ will be 185 m*g*. Another child may accumulate 60 min with accelerations just exceeding 200 m*g*. With the cut-point method, these similar activity levels look very disparate; zero min of MVPA and 60 min of MVPA, respectively. Their M60_ACC_, on the other hand reflects the smaller discrepancy in activity level that is evident; 185 m*g* and 200 m*g*. At a group level, presenting percentiles for the MX_ACC_ values as illustrated herein (Figures 1 and 2), displays the proportion of a sample achieving X min at any given intensity. In contrast, once cut-points have been applied, any activity accumulated just below a given cut-point will always be disregarded, irrespective of how the data are presented.

By decreasing the number of minutes of interest the metric can be used to focus on aspects of health that benefit from short, high-intensity bursts of activity, e.g. bone health^16, 29^. Accelerometer-derived physical activity intensity guides for bone health have recently been proposed for pre- and post-menopausal women using data from a UK Biobank^16^; these metrics could be used to further test this recommendation and to derive guidelines from accelerometer data specific to bone health in men and children.

To aid translation, we expressed the acceleration magnitudes in relation to indicative activities, e.g. brisk walk, fast walk and run. Currently there are limited data from which to draw these estimates. To enhance translation of these metrics there is a need to generate more data showing the acceleration ranges associated with indicative activities across a wide range of demographics. Note, this is only for translation and is not necessary for generation of the accelerometer metrics from data, or for developing the evidence base necessary to derive physical activity guidelines directly from accelerometer data.

The acceleration magnitudes tended to be higher for the pre-menopausal women who wore the Axivity on their dominant wrist than for the slightly younger office workers who wore the ActiGraph on their non-dominant wrist. While this may be due to the sedentary nature of the office job, it could reflect the non-representative nature of the samples, indicate that the −10% reduction in acceleration for dominant wrist placement^24^ was insufficient, and/or that there were differences between the ActiGraph and the Axivity. While raw data from the GENEActiv and Axivity accelerometers compare well^24^, ‘raw’ data from the ActiGraph GT9X is passed through a filter that suppresses higher intensity accelerations. The accelerometer sampling frequency and epoch differed between some studies. As the metrics are sampling frequency independent this should not impact on the outcomes generated with GGIR, but this needs to be confirmed empirically. It is also possible that the use of 1 s and 5 s epochs may have impacted on the MX_ACC_ outcomes, however, in our previous study data summarised in 1 s and 5 s epochs were comparable^30^.

## Conclusion

Cut-point approaches to analysing accelerometer data are not appropriate for assessing the prevalence of meeting guidelines globally^11^. Metrics reflecting the acceleration above which the most active minutes are accumulated are a standardised, easy to interpret, and population-independent method appropriate for assessing prevalence of physical activity and comparing activity between demographics and/or studies. These simple to derive variables facilitate global surveillance and dose-response studies. Furthermore, translating the metrics in terms of indicative activities (e.g. brisk walking) can provide a public-health friendly interpretation of the results^9^. Currently, guidelines are largely derived from self-report data^10^. As accelerometer and corresponding health data accumulate it will be possible to derive evidence-based physical activity guidelines directly from accelerometer data.

## Acknowledgements

The authors thank project staff involved with ASSK and Dr Sarah Taylor for data collection; all researchers and project staff involved in the Girls Active evaluation, SMArT Work trial, UK Biobank and CODEC (adults with type 2 diabetes) for access to the data used herein. Analysis of the pre-menopausal and post-menopausal samples was conducted using the UK Biobank Resource (Reference 10995).

We also thank: the participating schools in ASSK, children and teachers for their participation; the pupils and teachers who took part in the Girls Active evaluation study, the Youth Sport Trust (YST); the participants in the SMArT Work trial; the participants in UK Biobank; and participants in the CODEC study.

## References

1. da Silva ICM, van Hees VT, Ramires VV et al. Physical activity levels in three Brazilian birth cohorts as assessed with raw triaxial wrist accelerometry. Int J Epidemiol. 2014; 43(6):1959–1968.

2. Doherty A, Jackson D, Hammerla N et al. Large Scale Population Assessment of Physical Activity Using Wrist Worn Accelerometers: The UK Biobank Study. PLoS ONE. 2017; 12(2):e0169649. doi.org/10.1371/journal.pone.0169649.

3. Freedson PS, John D. Comment on “Estimating activity and sedentary behaviour from an accelerometer on the hip and wrist.” Med Sci Sports Exerc. 2013; 45(5):962–3.

4. Li X, Kearney PM, Keane E et al. Levels and sociodemographic correlates of accelerometer-based physical activity in Irish children: a cross-sectional study. J Epidemiol Community Health. 2017; 71(6):521–527. doi: 10.1136/jech-2016-207691.

5. Menai M, van Hees VT, Elbaz A et al. Accelerometer assessed moderate-to-vigorous physical activity and successful ageing: results from the Whitehall II study. Sci Rep 2017; 8:45772. doi:10.1038/srep45772.

6. World Health Organisation (2018). Global action plan on physical activity 2018–2030: more active people for a healthier world. Geneva:World Health Organization; 2018. Licence: CC BY-NC-SA 3.0 IGO. 104 p. Available from http://apps.who.int/iris/bitstream/handle/10665/272722/9789241514187-eng.pdf. Accessed 25th November 2018.

7. White T, Westgate K, Wareham NJ et al. Estimation of Physical Activity Energy Expenditure during Free-Living from Wrist Accelerometry in UK Adults. PLoS ONE 2016; 11(12):e0167472. doi:10.1371/journal.pone.0167472

8. Brazendale K, Beets MW, Bornstein DB et al. Equating accelerometer estimates among youth: The Rosetta Stone 2. J Sci Med Sport. 2016; 19:242–249.

9. Rowlands AV. Moving forward with accelerometer-assessed physical activity: Two strategies to ensure Meaningful, Interpretable & Comparable measures. Pediatr Exerc Sci. 2018; 30:450–456. doi.org/10.1123/pes.2018-0201.

10. Troiano RP, McClain JJ, Brychta RJ et al. Evolution of accelerometer methods for physical activity research. Brit J Sports Med. 2014; 48:1019–1023.

11. Migueles JH, Cadenas-Sanchez C, Tudor-Locke C et al. Comparability of published cut-points for the assessment of physical activity: Implications for data harmonization. Scan J Med Sci Sports. 2019; 1–9. doi.org/10.1111/sms.13356.

12. Physical Activity Guidelines for Americans 2018. 2nd edition. Washington, DC: US Department of Health and Human Services, 2018. 118 p. Available from: https://health.gov/paguidelines/secondedition/pdf/Physical_Activity_Guidelines_2nd_edition.pdf. Accessed November 18th, 2018.

13. Tudor-Locke C, Han H, Aguiar EJ et al. How fast is fast enough? Walking cadence (steps/min) as a practical estimate of intensity in adults: a narrative review. Brit J Sports Med 2018; 52(12):776–788. doi.org/10.1136/bjsports-2017-097628

14. Hildebrand M, Van Hees VT, Hansen BH. Age-group comparability of raw accelerometer output from wrist- and hip-worn monitors. Med Sci Sport Exerc 2014; 46: 1816–1824.

15. Phillips LRS, Parfitt CG, Rowlands AV. Calibration of the GENEA accelerometer for assessment of physical activity intensity in children. J Sci Med Sport. 2013; 6:124–128.

16. Stiles VS, Metcalf BS, Knapp KM et al. A small amount of precisely measured high intensity habitual physical activity predicts bone health in pre- and post-menopausal women in UK Biobank. Int J Epidemiol. 2017; 46:1847–1856. doi: 10.1093/ije/dyx080.

17. Taylor S, Noonan R, Knowles Z et al. Evaluation of a Pilot School-Based Physical Activity Clustered Randomised Controlled Trial—Active Schools: Skelmersdale. Int J Environ Res Public Health. 2018; 15(5):1011.

18. Edwardson CL, Harrington DM, Yates T et al. A cluster randomised controlled trial to investigate the effectiveness and cost effectiveness of the ‘Girls Active’ intervention: a study protocol. BMC Public Health. 2015; 15(1):526.

19. Harrington D, Davies MJ, Bodicoat DH et al. Effectiveness of the ‘Girls Active’ school-based physical activity programme: A cluster randomised controlled trial. Int J Behav Nutr Phys Act. 2018; 15:40. doi.org/10.1186/s12966-018-0664-6

20. Edwardson CL, Yates T, Biddle SJH et al. Effectiveness of the Stand More AT (SMArT) Work intervention: cluster randomised controlled trial. BMJ. 2018; 363:k3870. doi.org/10.1136/bmj.k3870.

21. van Hees VT, Fang Z, Langford J et al. Auto-calibration of accelerometer data for free-living physical activity assessment using local gravity and temperature: an evaluation on four continents. J Appl Physiol. 2014; 117(7): 738–744.

22. van Hees VT, Gorzelniak L, Dean León EC et al. Separating Movement and Gravity Components in an Acceleration Signal and Implications for the Assessment of Human Daily Physical Activity. PLoS ONE. 2013; 8(4):e61691. doi: 10.1371/journal.pone.0061691.

23. Rowlands AV, Edwardson CL, Davies MJ et al. Beyond cut-points: Accelerometer metrics that capture the physical activity profile. Med Sci Sport Exerc. 2018; 50(6):1323–1332.

24. Rowlands AV, Plekhanova T, Yates T et al. A basis for harmonisation of accelerometer physical activity outcomes in epidemiology. J Measure Phys Behav. 2019. In press.

25. Ainsworth BE, Haskall WL, Whitt MC et al. Compendium of physical activities: an update of activity codes and MET intensities. Med Sci Sport Exerc. 2000; 32:S498–504.

26. Lyden K, Keadle SK, Staudenmeyer J et al. Energy Cost of Common Activities in Children and Adolescents. J Phys Act Health. 2013; 10:62–69.

27. Cole TJ, Freeman JV, Preece MA. Body mass index reference curves for the UK, 1990. Arch Dis Child. 1995; 73:25–29.

28. Welk GJ, Bai Y, Lee J-M et al. Standardizing analytic methods and reporting in activity monitor validation studies: Guidelines to advance research and practice. Med Sci Sport Exerc. 2019. In press

29. Turner CH, Robling AG. Designing exercise regimens to increase bone strength. Exerc Sport Sci Rev. 2003; 31:45–50.

30. Rowlands AV, Cliff DP, Fairclough SJ et al. Moving forward with backwards compatibility: Translating wrist accelerometer data. Med Sci Sport Exerc. 2016; 48:2142–2149. doi: 10.1249/MSS.0000000000001015

